# *Staphylococcus aureus* COL: An Atypical Model Strain of MRSA that Exhibits Slow Growth and Antibiotic Tolerance Due to a Mutation in PRPP Synthetase

**DOI:** 10.1101/2024.12.12.627954

**Authors:** Claire E. Stevens, Ashley T. Deventer, Paul R. Johnston, Phillip T. Lowe, Alisdair B. Boraston, Joanne K. Hobbs

## Abstract

Methicillin-resistant *Staphylococcus aureus* (MRSA) has been a pathogen of global concern since its emergence in the 1960s. As one of the first MRSA strains isolated, COL has become a common model strain of *S. aureus*. Here we report that COL is, in fact, an atypical strain of MRSA that exhibits slow growth and multidrug tolerance. Genomic analysis identified three mutated genes in COL (*rpoB, gltX* and *prs*) with links to tolerance. Allele swapping experiments between COL and the closely related, non-tolerant Newman strain uncovered a complex interplay between these genes. However, Prs (phosphoribosyl pyrophosphate [PRPP] synthetase) accounted for most of the growth and tolerance phenotype of COL. Biochemical and transcriptomic analysis revealed that COL does not exhibit slow growth as a result of partial stringent response activation, as previously proposed. Instead, the COL Prs mutation greatly reduces the PRPP synthetase activity of the enzyme and leads to downregulation of pyrimidine, histidine and tryptophan synthesis, three pathways that rely on PRPP. Overall, our findings indicate that COL is an atypical, antibiotic-tolerant strain of MRSA whose isolation predates the previous first report of tolerance among clinical isolates. Characterisation of clinical Prs mutations and their relationship with tolerance requires further investigation.

## Introduction

*Staphylococcus aureus* is an opportunistic pathogen and major cause of both nosocomial and community-acquired infections. It is a leading cause of many life-threatening infections, including bacteremia, pneumonia, endocarditis, and bone and joint infections (Tong et al., 2015). Treatment of *S. aureus* infections is highly dependent on antibiotics, but *S. aureus* has an incredible capacity to develop resistance to these agents. The most widespread and clinically significant resistance occurs in methicillin-resistant *S. aureus* (MRSA). Methicillin resistance first emerged in the United Kingdom in 1960, within one year of the introduction of methicillin into clinical use (although genomic evidence suggests that MRSA actually first emerged in the 1940s (Harkins et al., 2017)). Since then, MRSA has proved itself to be a formidable global pathogen, causing at least 100,000 deaths in 2019, more than any other single pathogen-drug combination (Murray et al., 2022).

Methicillin resistance in *S. aureus* is most commonly mediated by *mecA*, which is acquired horizontally as part of a mobile genetic element known as staphylococcal cassette chromosome *mec* (SCC*mec*) (Miragaia, 2018). The *mecA* gene itself encodes for penicillin-binding protein 2a (PBP2a), which participates in peptidoglycan synthesis and has a lower binding affinity for β-lactam antibiotics than the native PBPs produced by *S. aureus*. Expression of methicillin resistance is multifactorial and supported by the presence and/or mutation of many “auxiliary” and “potentiator” genes (in addition to *mecA*) (Bilyk et al., 2022). Most clinical isolates of MRSA exhibit low-level and heterogeneous resistance; the population, as a whole, exhibits a relatively low MIC, but subpopulations of cells exhibit very high levels of resistance (Tomasz et al., 1991).

However, some isolates – like the early MRSA isolate COL – exhibit high-level, homogeneous resistance (De Lencastre et al., 1999). COL (previously known as strain 9204 and MR-COL in early literature) was one of the first MRSA strains identified, isolated in 1960 in a public health laboratory in Colindale, UK (Dyke et al., 1966; Sabath et al., 1972; Wilkinson et al., 1978). As a member of the so-called “archaic” lineage of MRSA (Bowers et al., 2018; Chambers and DeLeo, 2009), COL has become one of the most commonly used model strains of MRSA (*e.g.* Goetz et al., 2022; Lama et al., 2012; Madrigal et al., 2005; Reed et al., 2015; Surewaard et al., 2016; Tattevin et al., 2010; Tuffs et al., 2022; Xiao et al., 2014; Yeo et al., 2021).

COL is a member of clonal complex 8 (CC8), one of the most prevalent complexes of *S. aureus* responsible for both community-acquired and healthcare-associated infections (Bowers et al., 2018). Other notable model strains belonging to CC8 include the methicillin-sensitive *S. aureus* (MSSA) strain Newman and MRSA strains of the USA300 lineage (Table 1). COL is commonly employed as a model strain of *S. aureus* (*e.g.* Goetz et al., 2022; Lama et al., 2012; Madrigal et al., 2005; Reed et al., 2015; Surewaard et al., 2016; Tattevin et al., 2010; Tuffs et al., 2022; Xiao et al., 2014; Yeo et al., 2021), but two studies have noted that it exhibits slow growth compared with other CC8 strains (Kim et al., 2017; Li et al., 2009). Li et al. determined growth curves for COL and five comparator CC8 strains *in vitro* and noted that, while the growth curves of the other strains were virtually indistinguishable, COL exhibited “slightly slower growth” (Li et al., 2009). Later, Kim et al. also reported that COL exhibited slow growth but, more interestingly, they reported that COL maintained a higher basal concentration of the signalling molecules guanosine tetra- and pentaphosphate (collectively known as [p]ppGpp) than a comparator strain (Kim et al., 2017). (p)ppGpp is the effector molecule of the stringent response, a universal bacterial stress response that is classically induced in response to amino acid starvation (Irving et al., 2021). In *S. aureus*, activation of the stringent response and cellular (p)ppGpp concentration is largely controlled by a bifunctional enzyme, Rel, that both synthesises and hydrolyses (p)ppGpp (Atkinson et al., 2011). Once synthesised, (p)ppGpp acts to downregulate most metabolic processes through a range of direct and indirect mechanisms in different bacteria (Hauryliuk et al., 2015). In *S. aureus* and other members of the Firmicutes, much of the regulation of transcription and translation mediated by (p)ppGpp is thought to occur indirectly as a result of the plummeting intracellular GTP level (Rel synthesises (p)ppGpp from GTP). The transcriptional repressor CodY requires GTP as a ligand to facilitate binding to target DNA. Therefore, a rapid decrease in the intracellular GTP pool causes CodY to be released from DNA, allowing transcription of genes associated with, for example, amino acid synthesis (Geiger and Wolz, 2014). In contrast, the transcription of other genes is downregulated by the drop in GTP level, as many promoters (including those for rRNA genes) initiate with GTP (Krásný et al., 2008). Direct binding partners of (p)ppGpp have also been identified in *S. aureus* and closely-related bacteria, including enzymes involved in GTP synthesis and GTPases that participate in ribosome assembly (Corrigan et al., 2016; Kriel et al., 2012; Salzer and Wolz, 2023; Steinchen et al., 2020). Binding of (p)ppGpp to these protein targets inhibits their normal function, thereby contributing to the downregulation of metabolic processes.

**Table 1.** Model strains of *S. aureus* used in this study and MIC values.

| Strain name | Lineage | Clonal complex | Sequence type | Description | Genome accession number | Reference | MIC (mg/L)<sup>a</sup> |  |
| --- | --- | --- | --- | --- | --- | --- | --- | --- |
|  |  |  |  |  |  |  | Ciprofloxacin | Daptomycin |
| COL | Archaic | CC8 | ST250 | HA-MRSA | NC_002951 | (Dyke et al., 1966) | 0.125 | 0.5 |
| Newman | Archaic | CC8 | ST8 | MSSA | NC_009641 | (Duthie and Lorenz, 1952) | 0.125 | 0.5 |
| LAC | USA300 | CC8 | ST8 | CA-MRSA | CP035369 | (Voyich et al., 2005) | >8 | 0.5 |
| MW2 | USA400 | CC1 | ST1 | CA-MRSA | NC_003923 | (Baba et al., 2002) | 0.125 | 1 |
| N315 | - | CC5 | ST5 | HA-MRSA | BA000018 | (Kuroda et al., 2001) | 0.25 | 1 |
| MN8 | USA200 | CC30 | ST30 | MSSA | CP091875 | (Schlievert and Blomster, 1983) | 0.25 | 0.5 |

<sup>a</sup>MIC values were unchanged between wildtype and mutant variants of COL and Newman
Abbreviations: hospital-acquired (HA), community-acquired (CA), methicillin-sensitive *S. aureus* (MSSA).

We have previously shown that elevated cellular (p)ppGpp and a partially activated stringent response (as a result of clinical mutations in Rel) confers antibiotic tolerance in *S. aureus* Newman and USA300 strain LAC (Bryson et al., 2020; Deventer et al., 2022). Antibiotic tolerance describes the ability of a bacterial population to withstand transient exposure to an otherwise lethal concentration of bactericidal antibiotic, without exhibiting an elevated minimum inhibitory concentration (MIC) (Brauner et al., 2016). As a bacterial phenomenon, genotypic tolerance was first described in the pneumococcus in 1970 (Tomasz et al., 1970) and identified among clinical isolates of *S. aureus* in 1977 (Sabath et al., 1977). It has since been detected in >20 bacterial species and, arguably most frequently, in *S. aureus* (Deventer et al., 2024). In recent years, tolerance has been strongly implicated as a contributing factor in persistent and recurrent infections (Deventer et al., 2024; Kuehl et al., 2020; Lazarovits et al., 2022) and shown to promote the development of endogenous resistance (Levin-Reisman et al., 2017; Liu et al., 2020; Santi et al., 2021). While stringent response activation represents one route to tolerance, numerous other molecular mechanisms have been reported ranging from mutations in proteins associated with transcription, translation, carbon metabolism, purine biosynthesis and the electron transport chain (Deventer et al., 2024). An overarching theme that links most of these mechanisms is slow growth and reduced metabolism (Bryson et al., 2020; Deventer et al., 2024; Hobbs and Boraston, 2019; Proctor et al., 2014; Stokes et al., 2019). Most bactericidal antibiotics target active metabolic/cellular processes; therefore, slowing these processes down leads to a slower rate of cell death (Bren et al., 2023; Lee et al., 2018; Lobritz et al., 2015; Lopatkin et al., 2019; Tuomanen et al., 1986).

Given the reported slow growth phenotype of *S. aureus* COL (Kim et al., 2017; Li et al., 2009), we hypothesised that this property may confer antibiotic tolerance (as in other strains (Bryson et al., 2020; Deventer et al., 2024, 2022; Levin-Reisman et al., 2017; Liu et al., 2020)). Because COL is commonly used as a model strain of MRSA, it is important to understand any underlying mechanisms potentially impacting its behaviour in *in vitro* and *in vivo* antibiotic efficacy studies (Goetz et al., 2022; Lama et al., 2012; Madrigal et al., 2005; Surewaard et al., 2016; Tattevin et al., 2010; Xiao et al., 2014; Yeo et al., 2021). We, therefore, set out to investigate the tolerance profile of COL compared with other strains of *S. aureus.* Here, we show that COL exhibits a greatly extended lag phase and longer doubling time compared with other model strains of *S. aureus* and multidrug tolerance, but these phenotypes are not due to activation of the stringent response. Instead, genome analysis and allele swapping experiments demonstrate that the main contributor to the tolerance phenotype of COL is a mutation in Prs (phosphoribosyl pyrophosphate [PRPP] synthetase), a keystone enzyme in purine, pyrimidine, histidine, tryptophan and NAD^+^ biosynthesis. The COL Prs mutation causes a significant reduction in its catalytic activity and downregulation of three PRPP-utilising pathways. Overall, our results reveal that COL is an atypical strain of MRSA that exhibits slow growth and antibiotic tolerance primarily as a result of an uncommon mutation in Prs.

## Results

### COL exhibits growth defects and multidrug tolerance

In general, two types of tolerance have been identified: tolerance by lag and tolerance by slow growth (longer doubling time in exponential phase) (Bryson et al., 2020; Fridman et al., 2014). We have previously seen a combination of these two defects in a stringent response-activated mutant of *S. aureus* (Bryson et al., 2020). Li et al. reported that COL exhibited slower growth than other CC8 strains (Li et al., 2009). To investigate this further, we performed detailed growth curve analysis on COL and a panel of five other commonly used model strains of *S. aureus* from different CCs and sequence types (Table 1). While the mean lag times of the comparator strains didn’t differ significantly from each other (*P*>0.05; Figure 1), the lag time of COL was ∼50% longer than any other strain. Likewise, the mean doubling times of the comparator strains were similar, while the doubling time of COL was significantly longer. Viable counting of stationary phase cultures of each strain revealed that COL reached a similar or higher density of cells compared with the other strains (Figure S1); therefore, its extended lag phase is not due to the inoculation of fewer viable cells.

**Figure 1.**
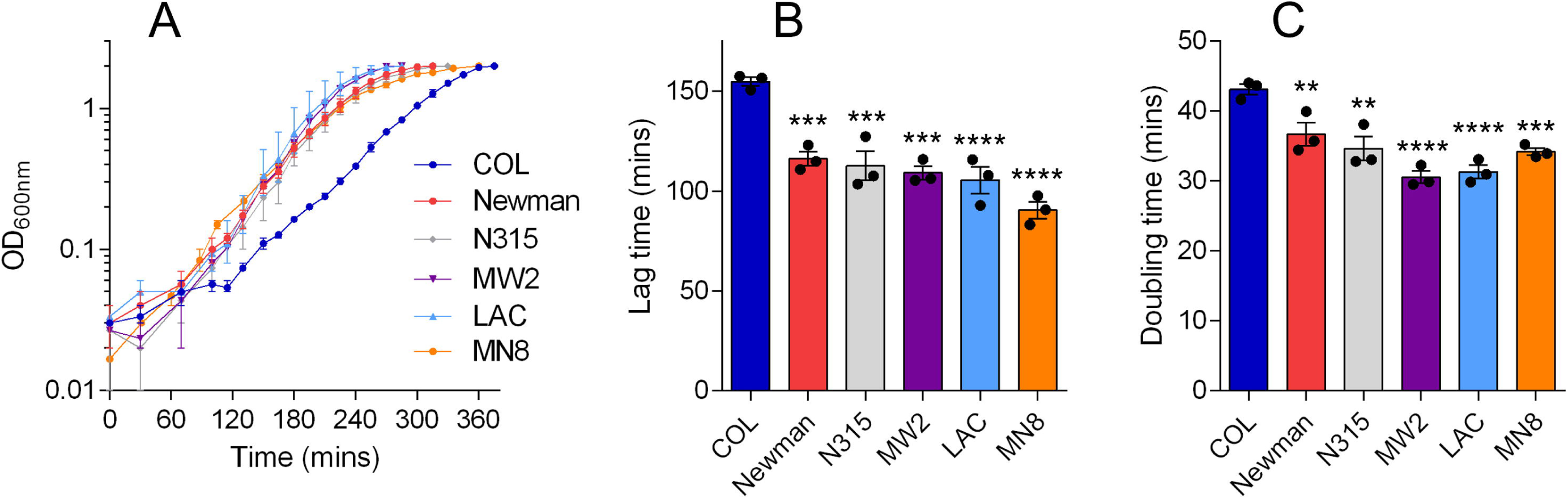
COL exhibits growth defects compared with other model strains of *S. aureus.* (A) Growth curves of six model *S. aureus* strains grown in TSB. Data shown are the mean of three biological replicates each derived from a different colony (error bars represent the range). Lag times (B) and doubling times (C) derived from the growth curves shown in panel A (error bars represent the SEM). Asterisks above bars indicate statically significant differences between means when compared with COL, as determined by a one-way ANOVA with Dunnett’s multiple comparisons test (**, *** and **** indicate *P* ≤0.01, *P* ≤0.001 and *P* ≤0.0001, respectively).

Antibiotic tolerance has not been reported previously for COL. However, given its growth defects, we hypothesised that it would exhibit this phenotype. We performed full time-kill assays with COL and comparator strains to compare their rates of antibiotic-induced death. We used two antibiotics with different mechanisms of action (at 8 or 16 × MIC; Table 1) and for which tolerance has been observed before in other strains of *S. aureus*: daptomycin and ciprofloxacin (Bryson et al., 2020; Corrigan et al., 2016; Liu et al., 2020). As predicted, the rate of killing of COL by daptomycin and ciprofloxacin was significantly slower than that of the comparator strains (Figure 2). The tolerance phenotype of COL to daptomycin was similar to that of our previously characterized tolerant *rel* mutant F128Y (Bryson et al., 2020; Deventer et al., 2022), while COL was more tolerant to ciprofloxacin than the *rel* mutant (Figure S2). We calculated the minimum duration for killing required to kill 99% of the population (MDK_99_), a metric used to quantify tolerance (Brauner et al., 2016), for each model strain-antibiotic combination. While there were some small but significant differences between the comparator strains (Figure 2C and D), the MDK_99_ values for COL were 50-300% higher than all other strains. These time-kill experiments were performed with cells that had been taken from stationary phase cultures and grown in fresh broth for 90 minutes prior to the addition of antibiotic. As the lag phase of COL is significantly longer than 90 minutes, the tolerance observed could be due to a difference in growth phase between COL and the other strains. As such, we repeated a time-kill with COL and Newman cultures that were grown to mid-exponential phase (OD_600nm_ ∼0.6) prior to the addition of antibiotic (Figure S3). Under these modified conditions, COL exhibited a similar level of tolerance to ciprofloxacin as determined with the previous experimental setup (∼200% increase in MDK_99_ compared with Newman; Figure S3). Therefore, irrespective of growth phase, COL appears to be an outlier with an atypical response to bactericidal antibiotics.

**Figure 2.**
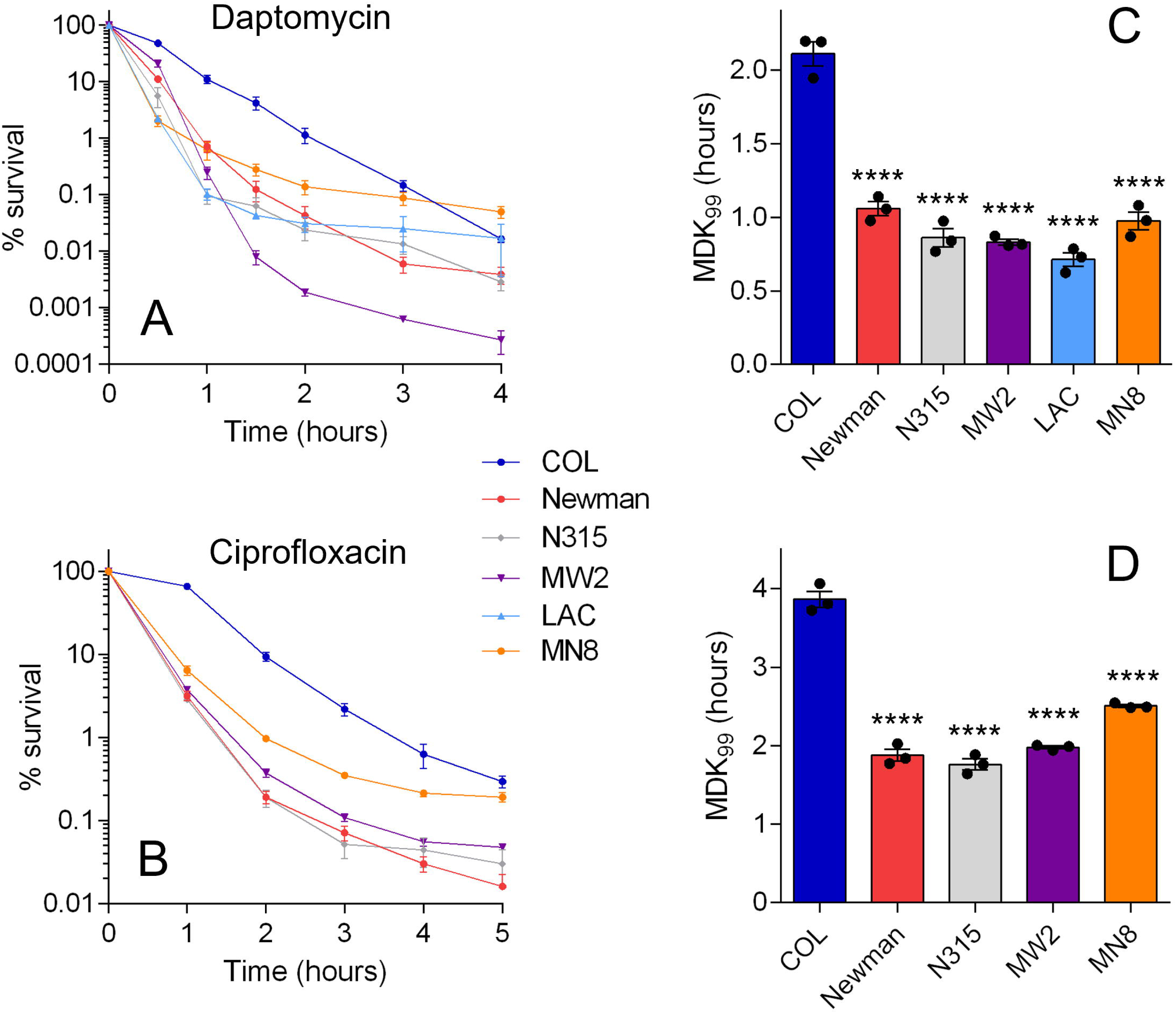
COL exhibits tolerance to daptomycin and ciprofloxacin compared with other model strains of *S. aureus.* Strains were exposed to (A) daptomycin or (B) ciprofloxacin and viable counts were determined at intervals. Counts at each time point are expressed as a percentage of the starting inoculum. Data shown are the mean of three biological replicates; error bars, where visible, represent the SEM. (C) and (D) Minimum duration for killing for 99% of the population (MDK_99_) values were interpolated from the data shown in panels A and B. Asterisks above bars indicate statically significant differences between means when compared with COL, as determined by a one-way ANOVA with Dunnett’s multiple comparisons test (**** indicates *P* ≤0.0001). Strain LAC is resistant to ciprofloxacin (Table 1) so is not included in panels B and D.

### Genome analysis of COL vs. comparator strains

Given the unusual growth and tolerance phenotype of COL compared to other strains of *S. aureus*, we were interested to investigate the genetic basis of these differences. In line with the fact that they belong to the same clonal complex, genome analysis revealed that COL is most similar to strains Newman and LAC, with 99.7% pairwise identity to both genomes. However, this still equates to >8,000 SNPs over a 2.8 kb genome. We have previously shown that a tolerance phenotype can be accompanied by a shift towards high-level homogeneous β-lactam resistance expression (Deventer et al., 2022), a phenotype that is also demonstrated by COL (De Lencastre et al., 1999). Therefore, in order to narrow down the pool of SNPs that may be contributing to the tolerance phenotype of COL, we considered previous work analysing the genetic basis of methicillin resistance expression in COL (De Lencastre et al., 1999). Kim et al. identified nonsynonymous mutations in four genes in COL that were deemed relevant to its high-level resistance expression: *prs* (ribose-phosphate pyrophosphokinase or phosphoribosyl pyrophosphate [PRPP] synthetase, involved in purine synthesis), *gltX* (glutamyl-tRNA synthetase), *rplK* (ribosomal protein L11) and *rpoB* (β-subunit of RNA polymerase) (Kim et al., 2017). A comparison of these protein sequences between COL and the five comparator strains revealed that the D103N mutation in *rplK* in COL is also found in Newman; therefore, it is unlikely to contribute to its tolerant phenotype. The mutations in *prs* (E112K), *gltX* (E405K) and *rpoB* (A798V and S875L), however, are exclusive to COL. Mutations in *prs*, *rpoB* and numerous tRNA-synthetases have been associated with tolerance previously (Deventer et al., 2024).

### Single allele swapping between COL and Newman

To experimentally test the impact of mutations in these three genes on growth and tolerance, we performed allele swapping between COL and Newman. We decided to use Newman, rather than LAC, as the comparator strain for these experiments because Newman is susceptible to ciprofloxacin which allowed us to perform time-kills with this antibiotic. We began by swapping the *prs* alleles between COL and Newman (Figure 3). Introduction of the COL *prs* allele into Newman lead to a dramatic growth defect, with a doubling time similar to that of wildtype COL and an even longer lag time. Introduction of the Newman *prs* allele into COL had no effect on doubling time, but it reduced the lag time to that of wildtype Newman. These effects were reflected in time-kills, where Newman::COL *prs* had a very similar response to ciprofloxacin as wildtype COL but was even more tolerant to killing by daptomycin than COL. With both antibiotics, the COL::Newman *prs* strain had a tolerant phenotype that was somewhere between that of wildtype Newman and COL. Interestingly, when exposed to daptomycin, COL::Newman *prs* exhibited a biphasic profile, with an initial rate of killing and MDK_99_ value indistinguishable from that of wildtype Newman (Figure 3E, Table 2) and a second, slower rate between that of Newman and COL. Collectively, these results indicate that the *prs* E112K mutation found in COL greatly contributes to its growth defects and tolerance phenotype.

**Figure 3.**
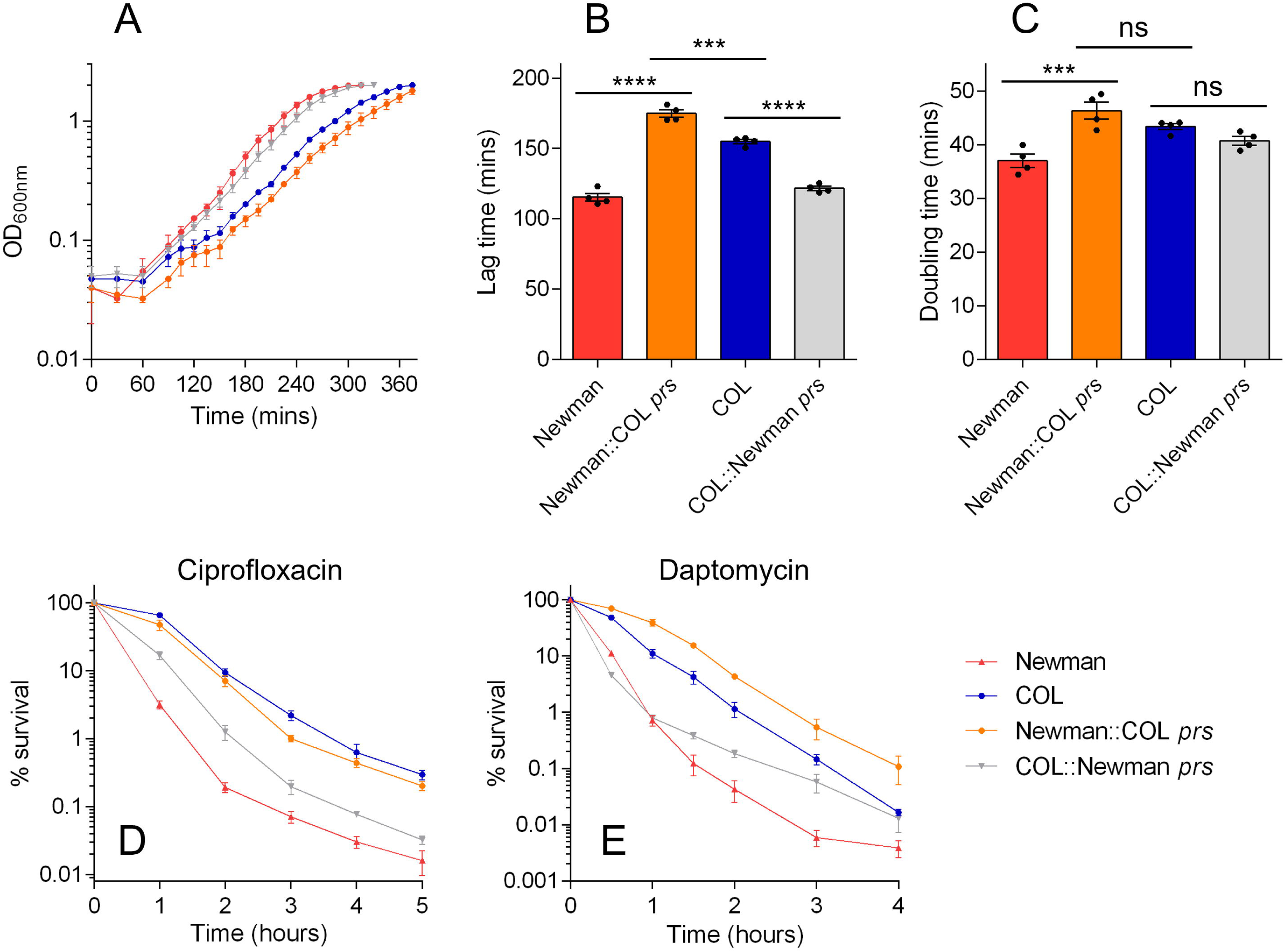
Impact of *prs* allele swapping on growth and tolerance. (A) Growth curves of wildtype COL, wildtype Newman and *prs* allele-swapped mutants grown in TSB. Data shown are the mean of four biological replicates each derived from a different colony (error bars represent the range). Lag times (B) and doubling times (C) derived from the growth curves shown in panel A. Asterisks above bars indicate statically significant differences between means when compared with COL, as determined by a one-way ANOVA with Dunnett’s multiple comparisons test (*** and **** indicate *P* ≤0.001 and *P* ≤0.0001, respectively; ns indicates not significant). (D) and (E) Time-kills with 4 × MIC ciprofloxacin and daptomycin, respectively. Data shown are the mean of three biological replicates; error bars (where visible) in panels B-E represent the SEM. See Table 2 for MDK_99_ values derived from data shown in panels D and E.

**Table 2.** Lag times, doubling times and MDK_99_ values for wildtype COL, wildtype Newman, and COL single, double and triple mutant strains*^a^*.

| Strain | Lag time (mins) | Doubling time (mins) | MDK<sub>99</sub> (CIP) | MDK<sub>99</sub> (DAP) |
| --- | --- | --- | --- | --- |
| COL | 155.7 ± 3.5 | 41.6 ± 1.7 | 3.9 ± 0.2 | 2.1 ± 0.1 |
| Newman | 115.3 ± 5.4<br>(74%****) | 37.0 ± 2.4<br>(89%**) | 1.9 ± 0.1<br>(49%****) | 1.1 ± 0.1<br>(52%**) |
| COL::Newman *prs* | 121.7 ± 3.0<br>(78%****) | 40.7 ± 1.6<br>(98%<sup>ns</sup>) | 2.5 ± 0.1<br>(64%****) | 1.2 ± 0.1<br>(57%*) |
| COL::Newman *gltX* | 129.8 ± 5.4<br>(83%****) | 44.2 ± 0.7<br>(106%<sup>ns</sup>) | 2.8 ± 0.4<br>(72%***) | 1.0 ± 0.1<br>(48%**) |
| COL *rpoB*<sup>+</sup> | 159.6 ± 3.6<br>(102%<sup>ns</sup>) | 48.7 ± 1.4<br>(117%***) | 3.7 ± 0.1<br>(95%<sup>ns</sup>) | 2.9 ± 0.1<br>(138%*) |
| COL::Newman *prs*::Newman *gltX* | 120.8 ± 3.8<br>(78%****) | 42.1 ± 1.3<br>(101%<sup>ns</sup>) | 2.0 ± 0.5<br>(51%****) | 1.7 ± 0.1<br>(81%<sup>ns</sup>) |
| COL *rpoB*<sup>+</sup>::Newman *prs* | 125.7 ± 3.6<br>(80.7%****) | 42.2 ± 2.0<br>(101%<sup>ns</sup>) | 1.3 ± 0.1<br>(33%****) | 0.9 ± 0.4<br>(43%***) |
| COL *rpoB*<sup>+</sup>::Newman *gltX* | 127.6 ± 2.0<br>(82%****) | 40.0 ± 2.9<br>(96%<sup>ns</sup>) | 1.6 ± 0.1<br>(41%****) | 1.7 ± 0.2<br>(81%<sup>ns</sup>) |
| COL *rpoB*<sup>+</sup>::Newman *prs*::Newman *gltX* | 110.3 ± 3.4<br>(71%****) | 35.8 ± 3.5<br>(86%**) | 2.4 ± 0.1<br>(62%****) | 1.4 ± 0.4<br>(67%**) |

<sup>a</sup>Percentages in parentheses indicate comparison with the value for COL. Asterisks indicate statically significant differences between means when compared with COL, as determined by a one-way ANOVA with Dunnett's multiple comparisons test (\*, \*\*, \*\*\* and \*\*\*\* indicate $P \leq 0.05$ , $P \leq 0.01$ , $P \leq 0.001$ and $P \leq 0.0001$ , respectively; ns indicates not significant).

Next, we moved on to the *gltX* allele and performed equivalent experiments. Introduction of the COL *gltX* allele into Newman had no effect on growth or tolerance to ciprofloxacin (Figure 4). The response of Newman::COL *gltX* to daptomycin was also very similar to that of wildtype Newman, although killing after two hours was reduced in the mutant. When the Newman *gltX* allele was introduced into COL, we observed no effect on doubling time and only a small (but significant) impact on lag time. However, this translated into a difference in antibiotic killing. Killing of COL::Newman *gltX* by ciprofloxacin was intermediate between that of wildtype COL and Newman, while the COL::Newman *gltX* strain exhibited a kill curve with daptomycin that was very similar to that of Newman. These results suggest that while the *gltX* mutation plays a role in tolerance in the genetic background of COL, this mutation alone is not sufficient to confer tolerance in Newman.

**Figure 4.**
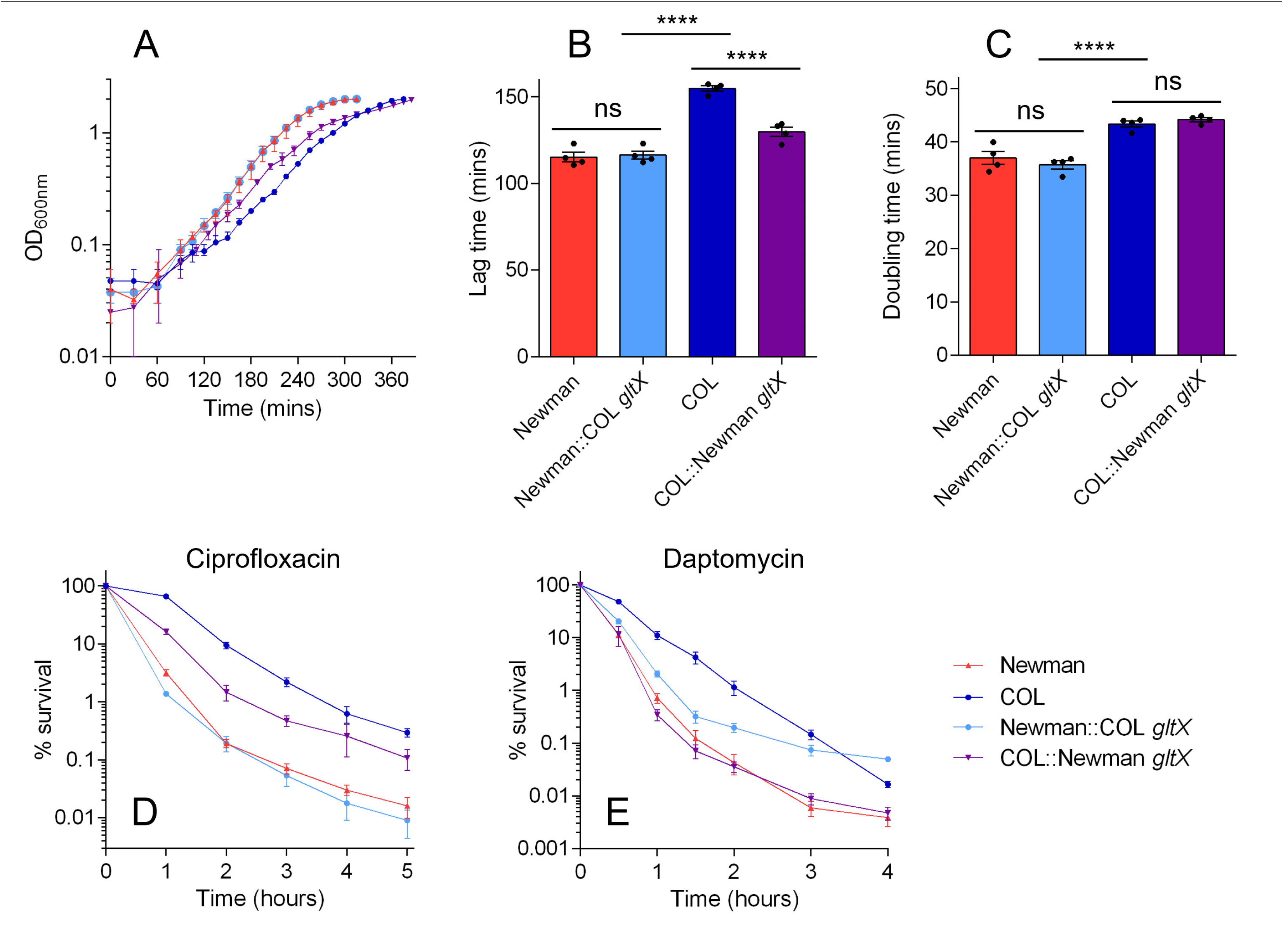
Impact of *gltX* allele swapping on growth and tolerance. (A) Growth curves of wildtype COL, wildtype Newman and *gltX* allele-swapped mutants grown in TSB. Data shown are the mean of four biological replicates each derived from a different colony (error bars represent the range). Lag times (B) and doubling times (C) derived from the growth curves shown in panel A. Asterisks above bars indicate statically significant differences between means when compared with COL, as determined by a one-way ANOVA with Dunnett’s multiple comparisons test (**** indicates *P* ≤0.0001; ns indicates not significant). (D) and (E) Time-kills with ciprofloxacin and daptomycin, respectively. Data shown are the mean of biological replicates; error bars (where visible) represent the SEM. See Table 2 for MDK_99_ values derived from data shown in panels D and E.

Finally, we wanted to assess the influence of the two *rpoB* mutations on the growth and tolerance of COL. Unfortunately, attempts to introduce the COL *rpoB* allele into Newman and vice versa were unsuccessful (likely due to the large size and essentiality of *rpoB*). However, Panchel et al. have previously generated COL *rpoB^+^*, a variant of COL in which the two *rpoB* mutations have been reversed in conjunction with the introduction of a nearby kanamycin resistance marker (Panchal et al., 2020). Therefore, we compared the growth and antibiotic killing profiles of wildtype COL and COL *rpoB^+^* (Figure 5). The lag times of the two strains were not significantly different, but, interestingly, the doubling time of COL *rpoB^+^* was slightly longer than that of wildtype COL. This small change in doubling time did not result in any difference in killing kinetics with ciprofloxacin, but COL *rpoB*^+^ did exhibit a greater MDK_99_ for daptomycin than wildtype COL (Table 2; *P* = 0.001, unpaired two-tailed *t*-test). Therefore, the two COL *rpoB* mutations present in COL *rpoB*^+^ do not confer tolerance and, in fact, the genetic engineering in this strain has increased tolerance to daptomycin slightly.

**Figure 5.**
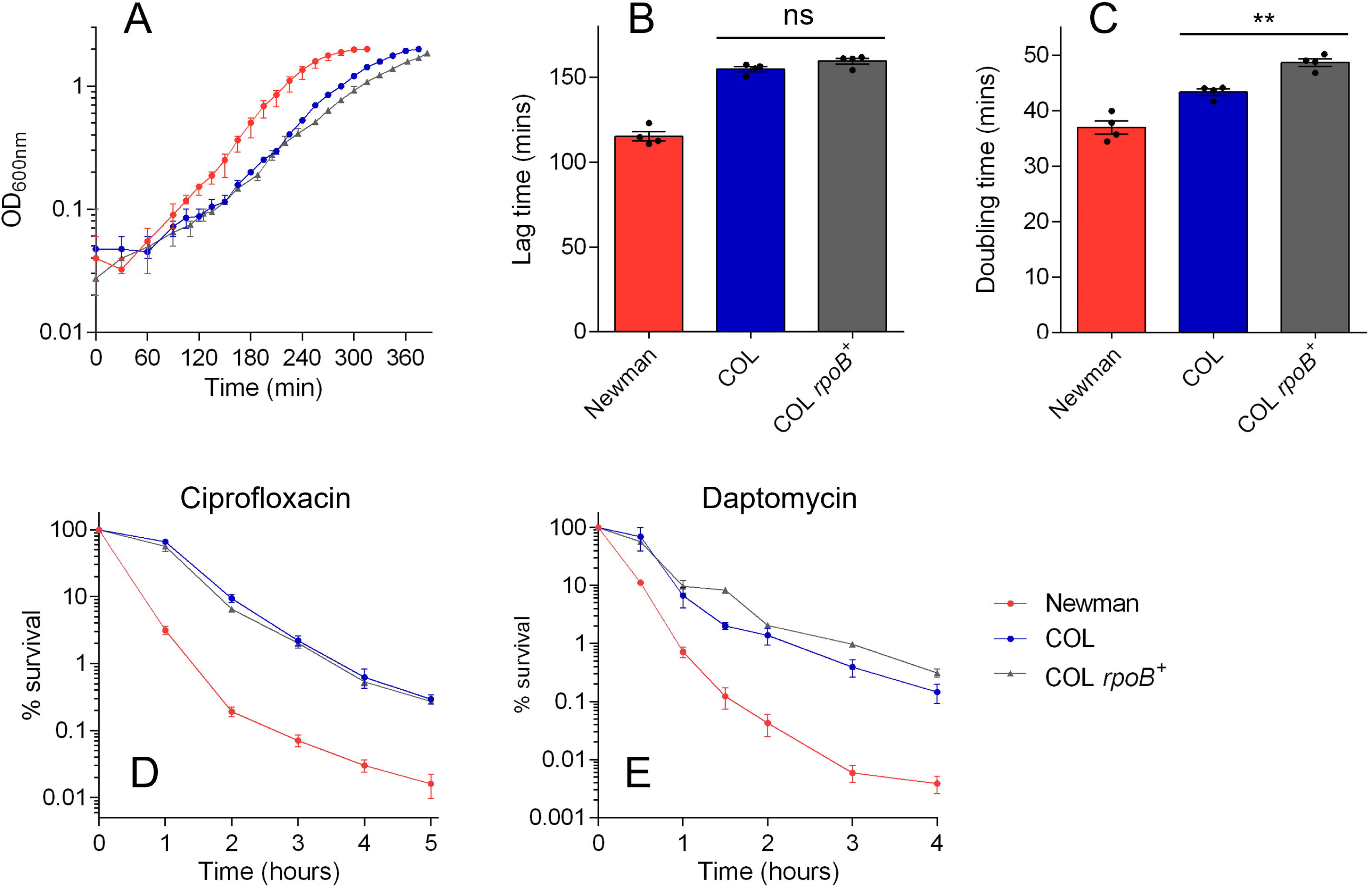
Impact of *rpoB* mutations on the growth and tolerance of COL. (A) Growth curves of wildtype COL, wildtype Newman and COL *rpoB*^+^ (a variant of COL that carries the Newman *rpoB* allele, as well as a kanamycin resistance gene) grown in TSB. Data shown are the mean of four biological replicates each derived from a different colony (error bars represent the range). Lag times (B) and doubling times (C) derived from the growth curves shown in panel A. Asterisks above bars indicate statically significant differences between means when compared with COL, as determined by a one-way ANOVA with Dunnett’s multiple comparisons test (** indicates *P* ≤0.01; ns indicates not significant). (D) and (E) Time-kills with ciprofloxacin and daptomycin, respectively. Data shown are the mean of three biological replicates; error bars (where visible) represent the SEM. See Table 2 for MDK_99_ values derived from data shown in panels D and E.

### Combining mutations in COL

Our single allele swapping experiments indicated that the mutation in *prs* is the major contributor to the reduced growth and tolerant phenotype of COL. However, reversing the *prs* mutation in COL did not fully abolish tolerance, and introduction of the Newman *gltX* allele into COL had a considerable effect on tolerance. Therefore, we decided to investigate the effect of multiple allele swapping in COL (Table 2, Figure S4). Replacing both the *prs* and *gltX* allelles in COL produced a strain with a lag time comparable to that of COL::Newman *prs* and a doubling time between that of COL::Newman *prs* and COL::Newman *gltX*. Its killing kinetics with both antibiotics were essentially the same as those of the *prs* single allele mutant; therefore, the effect of the *prs* allele appears to be more dominant than that of the *gltX* allele. When the *prs* allele was introduced into COL *rpoB*^+^, the lag time was indistinguishable from that of COL::Newman *prs* and the long doubling time of COL *rpoB*^+^ was reduced. There was no difference in the killing kinetics of COL::Newman *prs* and COL *rpoB*^+^::Newman *prs* with daptomycin, but the effect of replacing the *prs* allele on ciprofloxacin killing was greater in the presence of the *rpoB*^+^ allele than in wildtype COL (MDK_99_ of 1.3 h *vs* 2.5 h; Table 2). Therefore, there appears to be some combined effect of the *prs* and *rpoB* mutations. There also appears to be a complicated interaction between the *gltX* and *rpoB* alleles. Introduction of the Newman *gltX* allele into COL *rpoB*^+^ had the same effect on lag time that it did in wildtype COL, but it abolished the extended doubling time normally exhibited by COL *rpoB*^+^. In terms of tolerance, there appeared to be an additive effect of the *gltX* and *rpoB* mutations on tolerance to ciprofloxacin, as COL *rpoB*^+^::Newman *gltX* was killed more rapidly than COL::Newman *gltX*. However, with daptomycin, the reverse was observed and COL *rpoB*^+^::Newman *gltX* was more tolerant than COL::Newman *gltX*. Finally, when both the *prs* and *gltX* alleles were introduced into COL *rpoB*^+^, the resultant strain had the shortest lag time and doubling time of any COL mutant (Table 2). In terms of its tolerance profile, its killing kinetics were very similar to those of the COL *prs-gltX* double allele mutant, but it retained some tolerance to both ciprofloxacin and daptomycin compared with wildtype Newman.

### (p)ppGpp quantitation and stringent response transcriptomic signature

Our allelic exchange experiments suggest that the mutations in both Prs and GltX contribute to the tolerance phenotype of COL. Both enzymes have links with the stringent response; mutations in tRNA synthetases can limit the pool of charged tRNAs and induce (p)ppGpp synthesis (Vinella et al., 1992), while the purine biosynthesis pathway (of which Prs is a keystone enzyme) produces the substrates from which (p)ppGpp is synthesised (ATP and GDP/GTP). Furthermore, COL has previously been reported to exhibit higher (p)ppGpp content than a comparator strain (Kim et al., 2017). Therefore, we used (p)ppGpp quantitation and transcriptomics to compare the stringent response profiles of wildtype Newman and COL. Cellular extracts of exponentially-growing cultures of Newman, COL and a known stringent response-activated mutant of Newman that bears a Rel mutation, F128Y (Bryson et al., 2020), were subjected to HPLC separation and analysis using an established method for quantitation of ppGpp (the most abundant and stable stringent response alarmone (Varik et al., 2017)). While F128Y had a ppGpp level significantly greater than that of wildtype Newman and COL, the ppGpp content of COL did not differ significantly from that of Newman (Figure S5).

The lack of elevated ppGpp – the hallmark of stringent response activation – in COL was also corroborated by transcriptomic analysis of COL *vs* Newman. The hallmarks of stringent response activation in *S. aureus* are well known and include upregulation of 150 genes under regulation by CodY (Geiger et al., 2012). We identified 147 CodY regulon genes in our COL and Newman datasets. Of these 147 genes, 105 were not differentially expressed in COL compared with Newman, 30 were upregulated in COL and 12 were downregulated in COL (Figure S6, Table S1). Therefore, there is no transcriptomic evidence to support the hypothesis that the stringent response is induced in COL or that elevated (p)ppGpp contributes to its growth and tolerance phenotypes.

#### Molecular and cellular effects of Prs E112K mutation

The majority of the slow growth and tolerance phenotype of COL appears to be derived from the E112K mutation in Prs. Prs is a ribose-phosphate pyrophosphokinase that synthesises PRPP from ribose-5-phosphate and ATP, producing AMP as a byproduct. To investigate the impact of the E112K mutation on the catalytic activity of Prs, we produced recombinant wildtype Newman Prs and a E112K variant and compared their rates of activity in an enzyme-coupled assay that provides a readout of AMP. The mutant protein exhibited a >80% reduction in rate compared with the wildtype (Figure S7).

PRPP is a keystone metabolite that is essential not only for *de novo* purine synthesis, but also the purine salvage pathway and the synthesis of tryptophan, histidine, pyrimidines and NAD^+^ (Figure S8) (Hove-Jensen et al., 2017). Therefore, we explored the downstream cellular impact of reduced Prs activity by comparing the transcriptomic profiles of Newman *vs* Newman::COL Prs and COL *vs* COL::Newman Prs (Figure 6). Interestingly, these comparisons did not mirror each other as closely as we may have expected. If we focus in on the 40+ genes that lie downstream of *prs* in PRPP-utilising pathways, we see some interesting trends. Many of these genes are arranged in operons (ten Broeke-Smits et al., 2010) so, for simplicity, Figure 6A presents aggregated normalised counts for seven operons, as well as counts for a further 11 individual genes (data for individual genes is given in Tables S2 and S3). The 11-step pathway that converts PRPP into inosine monophosphate (IMP) requires 10 *pur* enzymes encoded in a single operon, plus *purB* (Figure 6A and Figure S8). Introduction of Prs E112K into Newman resulted in a strong upregulation of these 11 genes, while reversal of the Prs mutation in COL conferred a small, but significant, decrease in expression. The same trend was seen for PurA which, together with PurB, converts IMP into AMP. These results are consistent with the differential expression of *purR* (the transcriptional repressor of the *pur* operon and *purA* (Goncheva et al., 2019)) seen upon Prs mutation (Figure 6A). However, in the GTP branch of the *de novo* pathway, an unexpected profile was observed, whereby *guaA* and *guaB* expression was downregulated in both Newman::COL Prs *and* COL::Newman Prs. Conversely, *ndk*, which encodes for the enzyme that converts both ADP and GDP into their triphosphate forms, was upregulated in both mutant strains. Therefore, the majority of the *de novo* purine synthesis pathway is upregulated in response to the Prs E112K mutation, but there are some exceptions and differences between the ATP and GTP branches.

**Figure 6.**
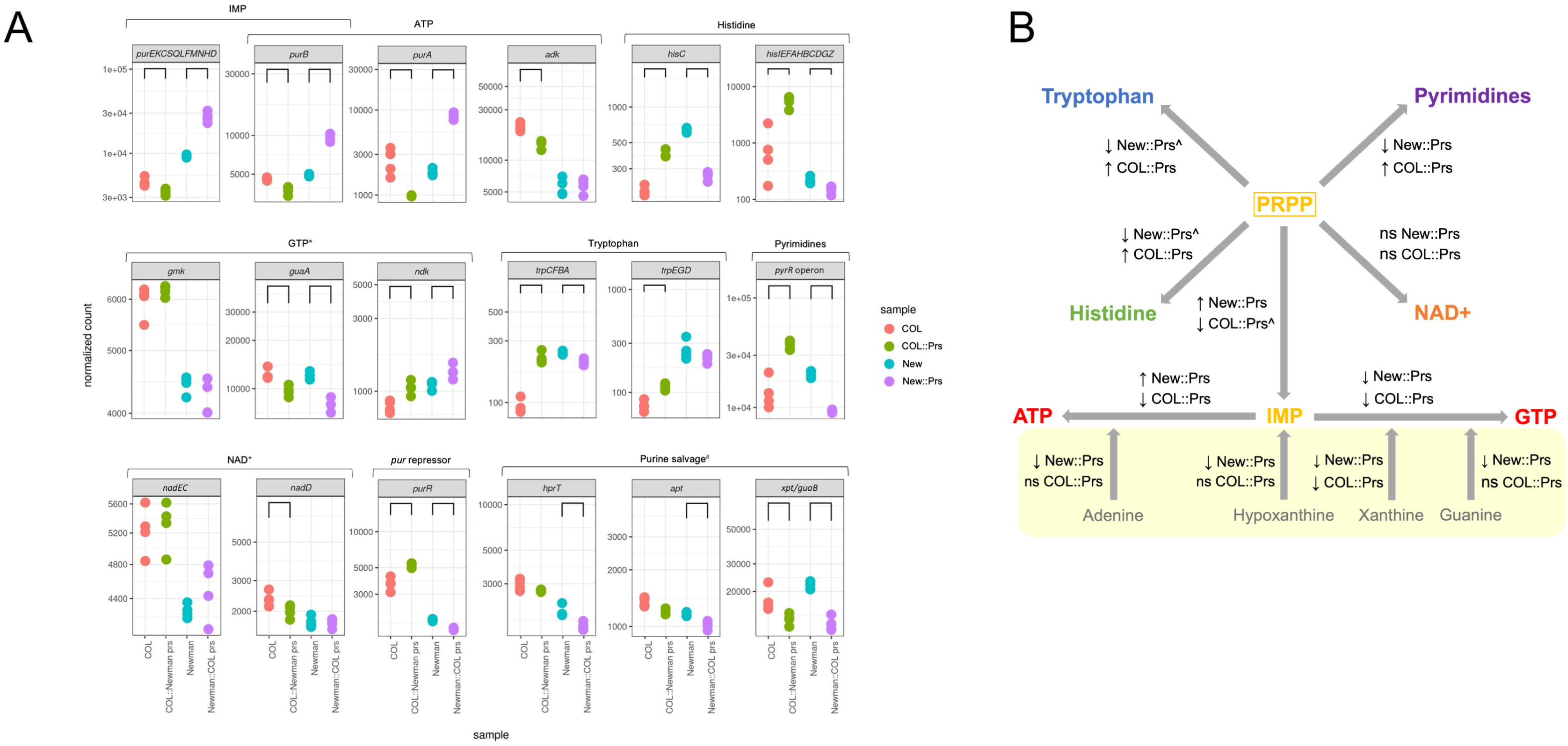
Impact of Prs mutation on PRPP-utilising pathway gene expression. (A) Normalised counts of operons and genes from metabolic pathways that use PRPP. Data for four biological replicates are shown. Brackets indicate counts that are significantly different between wildtype and mutant strains (FDR-adjusted *p*<0.05, no fold change threshold applied). Strain names have been abbreviated as follows: COL::Newman Prs (COL::Prs); Newman (New); Newman::COL Prs (New::Prs). A superscript × indicates that *ndk* is also involved in ATP synthesis, while superscript # indicates that *xpt* and *guaB* are in the same operon, but *guaB* is part of the *de novo* GTP pathway. (B) Summary of the overall differential gene expression trend for each PRPP-utilising pathway generated using the data shown in panel A. Small black arrows indicate the overall direction of differential gene expression in the strain shown; ns indicates no significant differential expression for the pathway as a whole. ^ indicates that the fold change was <2. The salvage purine synthesis pathway is shaded in yellow. See Figure S8 for more information on each pathway.

In terms of the other pathways that use PRPP, gene expression in the histidine, tryptophan and pyrimidine pathways showed a consistent trend of downregulation upon mutation of Prs in Newman and upregulation upon reversal of the Prs mutation in COL (although the fold change was not always >2; Figure 6). The NAD^+^ pathway did not show a consistent trend in differential gene expression in either strain comparison, although this pathway is predicted to only use a very small proportion of the PRPP pool (1-2%) (Hove-Jensen, 1988; Jensen, 1983). The purine salvage pathway also uses PRPP (Figure S8), and the three genes involved (*apt, hprT* and *xpt*) were consistently downregulated in Newman::COL Prs compared with wildtype Newman. *apt* and *hprT* were not significantly differentially expressed in COL::Newman Prs compared with wildtype COL; while *xpt* was significantly upregulated in the COL mutant, it is in an operon with *guaB* from the *de novo* GTP pathway. The growth defect exhibited by Newman::COL Prs could not be rescued by supplementation with tryptophan (which is unstable and typically lacking from culture media (Sezonov et al., 2007)) or a combination of tryptophan, histidine, inosine, guanosine and adenosine (Figure S9).

#### Bioinformatic analysis of Prs

The E112K Prs allele accounts for a large proportion of the extended lag time and tolerance phenotype of COL, and Prs mutations have been associated with tolerance in *in vitro*-evolved strains of both *S. aureus* and *Escherichia coli* before (Fridman et al., 2014; Levin-Reisman et al., 2017; Sulaiman and Lam, 2021). Therefore, we set out to analyse the prevalence of Prs mutations in *S. aureus*. Bioinformatic analysis of the >110,000 *S. aureus* genomes in GenBank reveals that >99% are identical; just 10 entries carry the Prs E112K mutation and 8 of these are derivatives of COL. The remaining two strains are: RN8098 (accession GCF_018132125.1), a strain that seemingly originated from the Novick group and carries the SaPI3 pathogenicity island; and a strain simply named “USA100” (accession GCA_016916755.1) and isolated in 1960. The latter seems likely to be a clinical isolate, but the reference cited does not include an isolate with this name. If we look specifically at a collection of sequenced *S. aureus* clinical isolates (the British Society for Antimicrobial Chemotherapy [BSAC] Resistance Surveillance Programme collection, https://www.ebi.ac.uk/ena/browser/view/PRJEB2756) and more generally at mutations in Prs, again, Prs is highly conserved and just three strains contain missense mutations: V155E, S272P and A304V. Based on previous mutational and structural analysis of Prs from *E. coli* and *Bacillus subtilis* (which shares 77% amino acid identity with the *S. aureus* enzyme) (Eriksen et al., 2000; Zhou et al., 2019), neither E112 or any of these three newly identified mutated residues overlap with sites known to be important for catalysis, allosteric regulation or intramolecular interactions (Figure S10).

## Discussion

COL is an archaic MRSA isolate that is frequently used as a model strain in *in vitro* and *in vivo* studies of *S. aureus* (Goetz et al., 2022; Lama et al., 2012; Madrigal et al., 2005; Surewaard et al., 2016; Tattevin et al., 2010; Xiao et al., 2014; Yeo et al., 2021). It has previously been identified as exhibiting high-level, homogeneous methicillin resistance expression, which is in contrast to most MRSA strains that exhibit low-level, heterogeneous resistance (Tomasz et al., 1991). Here, we have confirmed two previous reports (Kim et al., 2017; Li et al., 2009) that COL exhibits growth defects compared with other strains of *S. aureus* and have further shown that these defects lead to antibiotic tolerance. The slow growth of COL has previously been explained by an elevated intracellular concentration of (p)ppGpp compared with a comparator strain (Kim et al., 2017). However, our ppGpp quantitation experiments suggest that COL contains a similar basal level of ppGpp to its genetically close comparator Newman. Further, the transcriptomic profile of COL does not show the hallmarks of stringent response activation. The extended lag phase of COL is more pronounced than its modestly slow growth during exponential phase when compared with other model *S. aureus* strains. This adaptation towards a longer lag phase is consistent with the experimental evolution of “tolerance by lag” seen *in vitro*, whereby bacterial populations evolve to have a lag phase that is optimised to the interval of antibiotic exposure (Fridman et al., 2014). However, our time-kill results with exponentially growing cultures indicate that the tolerance phenotype of COL is consistent irrespective of its growth phase. What remains to be tested (both for COL and more generally among tolerant strains) is whether *in vitro* tolerance is observed *in vivo,* where the bacteria are growing in a less uniform and predictable environment.

Our allele swapping experiments focused on three genes that bear mutations in COL and that have been previously associated with tolerance: *rpoB, gltX* and *prs.* In isolation, the *rpoB* mutations A798V and S875L do not play a significant role in the tolerance phenotype of COL as their reversal had no effect on antibiotic killing. However, there appeared to be a small additive effect of these mutations when combined with the mutant alleles of *prs* or *gltX* (although these experiments are potentially confounded by the use of the marked COL *rpoB*^+^ strain, which grows slower than wildtype COL). Similarly, there is clearly an interplay between the GltX E405K mutation and the wider genetic background of COL in terms of tolerance. While the GltX mutation alone was insufficient to confer tolerance in Newman, reversal of this mutation in COL impacted antibiotic killing, and reversal in COL *rpoB^+^* had an even greater impact. Testing of the different single, double and triple allele-swapped mutants of COL also revealed some differences in tolerance and killing between ciprofloxacin and daptomycin, highlighting the importance of testing more than one antibiotic.

Interestingly, the COL triple mutant, with all three mutant alleles replaced, still exhibited some low-level tolerance to both antibiotics compared with Newman. Therefore, while we could conclude that these three genes may not represent the full genetic complement of tolerance in COL, tolerance is a not an absolute property but a relative phenotype and is only detected by comparison with a non-tolerant strain (in this case Newman). It should also be noted here that while the *prs* and *gltX* mutations in COL have been linked to its high-level, homogeneous methicillin resistance phenotype (Kim et al., 2017), reversal of the two *rpoB* mutations alone is sufficient to abolish this phenotype (Panchal et al., 2020).

Of the three genes we looked at, the E112K Prs allele accounts for a large proportion of the extended lag time and tolerance phenotype of COL. Therefore, we investigated this mutation in more detail. Our enzymology data indicate that E112K greatly reduces the rate of Prs catalytic activity. While E112 has not previously been identified as a residue important for substrate binding, intramolecular interactions or allosteric regulation (Eriksen et al., 2000; Zhou et al., 2019), it is found directly adjacent to an allosteric binding site (Figure S10). More in-depth enzymology, allosteric binding site affinity analysis and structural studies will be required to determine the molecular details underpinning the effect of this mutation. In terms of the downstream effects of the mutation, the transcriptomic analysis of our Prs allele swapped mutants suggests that the E112K mutation causes upregulation of the core steps in the *de novo* purine synthesis pathway, while downregulating the other pathways that use PRPP. However, some genes were differentially expressed in the same direction in both Newman::COL Prs and COL::Newman Prs, like *guaA* and *guaB* which are part of the GTP branch of the purine pathway. Therefore, as we saw with our growth curve and time-kill experiments with the different allele swapped mutants, the differences between Newman and COL cannot be accounted for solely by Prs and the effect of a single mutation is often dependent on the wider genetic background.

To our knowledge, COL represents the first reported case of a Prs mutation in a clinical isolate. However, other Prs mutations have been reported in *in vitro*-evolved slow growing and tolerant strains of *S. aureus* and *E. coli* under antibiotic pressure (Fridman et al., 2014; Levin-Reisman et al., 2017; Sulaiman and Lam, 2021), and here we have identified three new Prs mutations among clinical isolates of *S. aureus* from the BSAC collection. Therefore, disruption of PRPP synthesis may not be an uncommon mechanism of genotypic tolerance. In general, there are conflicting data in the literature regarding the relationship between purine synthesis, tolerance and persistent infections (Li et al., 2018; Xiong et al., 2024; Yang et al., 2019) which requires further investigation. Metabolomics and metabolic flux analysis of Prs mutants would be an important next step in understanding the impact of reduced PRPP synthetase activity on each of the downstream pathways. In particular, it would be interesting to compare the GTP pool between the mutant strains as the gene expression data for the GTP branch show the same effect when the Prs mutation is introduced into Newman as when it is reversed in COL. If reduced PRPP synthetase activity leads to a reduction in the GTP pool, this could potentially phenocopy stringent response activation in the absence of elevated (p)ppGpp (although no transcriptomic signature of stringent response activation was observed in COL).

COL is a commonly used model strain of MRSA in both *in vitro* and *in vivo* studies of antibiotic efficacy (Goetz et al., 2022; Lama et al., 2012; Madrigal et al., 2005; Surewaard et al., 2016; Tattevin et al., 2010; Xiao et al., 2014; Yeo et al., 2021). Therefore, identification of its tolerant phenotype is highly relevant and, given that the Prs E112K mutation is extremely rare, indicates that COL is an atypical strain of MRSA. While COL represents an invaluable and clinically relevant example of tolerant *S. aureus* for much-needed wider studies of tolerance (*e.g.* animal models of antibiotic efficacy, impact of tolerance on development of resistance) (Deventer et al., 2024), its unusual growth and tolerance phenotypes should be considered when using it as a model MRSA strain in other studies. Overall, identification of tolerance in COL, which was isolated in the 1960s (Dyke et al., 1966), shows that antibiotic tolerance in this strain predates the seminal first report of tolerance among clinical isolates by more than 10 years (Sabath et al., 1977). It also indicates that clinical antibiotic tolerance occurred in *S. aureus* around the same time as, or even before, clinical methicillin resistance. The clinical prevalence and significance of antibiotic tolerance, as well as its ability to promote the development of endogenous resistance, remains a largely unexplored area of clinical microbiology that requires attention.

### Experimental Procedures

#### Antibiotics and reagents

X-Gal (5-bromo-4-chloro-3-indolyl-β-d-galactopyranoside) and chloramphenicol were purchased from Bio Basic Inc. (Markham, ON, Canada). Daptomycin and ppGpp were obtained from Toronto Research Chemicals (Toronto, ON, Canada) and Jena Bioscience (Jena, Germany), respectively. All other antibiotics, enzymes, and reagents, unless otherwise stated, were from MilliporeSigma (Gillingham, UK).

#### Bacterial strains, plasmids and growth conditions

*S. aureus* strains Newman and MN8 were gifts from Michael Murphy (University of British Columbia) and Mariya Goncheva (University of Victoria, British Columbia), respectively, while strains COL, N315, MW2 and LAC were gifts from Paul Kubes (University of Calgary). COL *rpoB^+^* was a gift from Simon Foster (University of Sheffield). The Newman *rel* F128Y mutant has been reported previously (Bryson et al., 2020). *Escherichia coli* Stellar (TaKaRa Bio USA, Mountain View, CA) was used for all cloning, while *E. coli* IM08B (Monk et al., 2015) was used to prepare methylated plasmid for transformation into Newman and COL. Plasmid pIMAY-Z (Monk et al., 2015) was a gift from Ian Monk (University of Melbourne, Australia). *S. aureus* strains were routinely cultured in tryptic soy broth/tryptic soy agar (TSB/TSA) at 37°C and stored long term at −80°C with 8% (vol/vol) glycerol.

#### Allelic exchange

Allelic exchange of *prs* (NWMN_RS02645, SACOL0544) and *gltX* (NWMN_RS02865, SACOL0574) was performed using pIMAY-Z as previously described (Bryson et al., 2020). For allelic exchange in Newman, both genes were amplified from Newman genomic DNA as two halves using CloneAmp HiFi PCR premix (TaKaRa Bio) and the COL mutation-containing primers listed in Table S4 (E112K for *prs* and E405K for *gltX*). The two halves were then joined together by overlap extension PCR and cloned into pIMAY-Z between the *Eco*RI and *Not*I sites using In-Fusion Snap Assembly mastermix (TaKaRa Bio). To generate the constructs for allelic exchange in COL, *prs* was amplified from COL genomic DNA as two halves with mutagenic primers (K112E) as above, while the Newman *gltX* pIMAY-Z construct was mutated to K405E *via* site-directed mutagenesis. Following construct confirmation by bidirectional sequencing and passage through IM08B for methylation, constructs were introduced into Newman by electroporation and COL by phage transduction with *ϕ*85(Olson, 2016). pIMAY-Z integration and excision was performed as previously described (Bryson et al., 2020). Primers that bind outside of the mutated gene were used to amplify the region for confirmatory bidirectional sequencing (Table S2).

#### Growth curve analysis

Growth curves for all strains were performed in triplicate or quadruplicate in 5 mL of TSB in 16 mm test tubes. Overnight cultures derived from different colonies were diluted (10 uL in 5 mL fresh TSB) and grown at 37°C with 200 rpm shaking. OD_600nm_ was recorded using a test tube spectrophotometer every 30 minutes for the first 90 minutes, and then every 15 minutes until an OD_600nm_ of 2.0 was reached. The spectrophotometer used had an experimentally determined linear range of 0.05-1.5 (R^2^ = 0.984). Specific growth rates (μ) were determined from the slope of ln(OD_600nm_) versus time during exponential phase (maximum OD_600nm_ values of 0.8), and converted into doubling times using the equation ln(2)/μ. It should be noted that these doubling times represent doubling of culture density as determined by OD_600nm_ and, as such, doubling of colony forming units (CFUs) cannot be inferred. Lag times were found by extrapolating the slope of the exponential phase back to an OD_600nm_ of 0.1. All statistical analysis was performed in GraphPad Prism 6.07. Supplementation growth assays were performed in a 96-well plate in a Tecan Spark plate reader at 37°C with continuous shaking and OD_600nm_ readings every 10 minutes. TSB was supplemented with 0.1 mM each of tryptophan, histidine, inosine and guanosine, and 0.4 mM adenosine as previously described (Koenigsknecht et al., 2010), aliquoted into wells (200 μL) and inoculated with 1 μL over overnight culture. Three biological replicates were performed per strain per condition.

#### Minimum inhibitory concentration (MIC) determinations

Strains of interest were grown overnight in 5 mL of Mueller-Hinton broth (MHB) at 37°C with 200 rpm shaking. MIC determinations were performed in triplicate using the broth microdilution method of the Clinical and Laboratory Standards Institute (CLSI, 2024) in MHB or MHB supplemented with 50 μg/mL Ca^2+^ (MHB + Ca^2+^) for daptomycin.

#### Time-kill assays

Standard time-kill assays were performed in triplicate in MHB or MHB + Ca^2+^ in 18 mm test tubes at 37°C according to CLSI guidelines (CLSI, 1999). Briefly, overnight cultures were diluted 1 in 500 and grown for 90 mins prior to the addition of antibiotic (the cell density at this point was ∼10^6^ CFU/mL). Antibiotics were tested at 8 × MIC for daptomycin and 16 × MIC for ciprofloxacin for all strains. Aliquots were taken from growing cultures at defined time points, diluted in phosphate-buffered saline and plated in duplicate on TSA agar. Colony forming units (CFU) were counted using the aCOLyte 3 HD automated colony counter (Synbiosis Ltd, Cambridge), with counts periodically confirmed manually. The limit of detection following antibiotic dilution was 300 CFU/mL. MDK_99_ values were calculated using linear regression and data interpolation in GraphPad Prism 6.07; only data in the linear range of the MDK were used in the calculation (a minimum of three data points) and all *R*^2^ values were >0.9. For the ciprofloxacin time-kill with exponentially growing cells, strains were grown to an OD_600nm_ of 0.6 and diluted to ∼10^6^ CFU/mL prior to the addition of antibiotic.

#### (p)ppGpp quantitation by HPLC

(p)ppGpp extraction and quantitation was performed largely as described by Varik et al., 2017. Quadruplicate cultures of 250 mL in TSB were inoculated with 5 mL of overnight culture (originating from different colonies), grown at 37°C with shaking to an OD_600nm_ of 0.6-0.8 and filtered through 45 µm cellulose acetate filters (6 per sample) using a vacuum manifold (MilliporeSigma). Filters were immediately added to a 15 mL falcon tube containing 5 mL 1 M acetic acid, then flash frozen in liquid nitrogen and stored at - 80°C. To extract nucleotides, samples were thawed on ice then vortexed for 5-10 seconds, every 5 minutes, for 30 minutes and kept on ice in between. Filters were removed and the extract flash frozen in liquid nitrogen and stored at −80°C. Samples were freeze-dried overnight, resuspended in 600 µL of milliQ water, centrifuged to remove cell debris, and the supernatant stored at −20°C until analysis. Nucleotides were separated *via* strong-anion exchange (SAX) chromatography using a SphereClone SAX (4.6 × 150 mm, 5 μm) column fitted with a SecurityGuard cartridge (Phenomenex Ltd, Macclesfield, UK) run on a Shimadzu HPLC system (UV detection: 252 nm) using a 32 minute isocratic program with 0.36 M NH_4_H_2_PO_4_ pH 3.4, 25 % (v/v) acetonitrile and a flow rate of 0.5 mL/min. Elution time and peaks corresponding to ppGpp were identified by spiking every 8^th^ run with 2 mM ppGpp (Jena Bioscience). Area under the peak for ppGpp was calculated using LabSolutions software (Shimadzu) and corrected for the OD_600nm_ of the culture at harvest.

#### RNA sequencing

RNA was extracted from 2 mL of quadruplicate cultures grown in TSB to OD_600nm_ ∼0.6. Cultures were combined with 2 volumes of RNAprotect® (Qiagen, Manchester, UK) and incubated for 5 mins at room temperature, then centrifuged at 5000 rpm for 10 mins. The supernatant was removed and frozen at −20°C until extraction. RNA was extracted using the RNeasy kit (Qiagen) according to the manufacturer’s instructions using both mechanical disruption and chemical disruption with lysostaphin. Following extraction, the samples were treated with DNase and incubated at 37°C for 30 mins to eliminate contamination with DNA. RNA was then repurified using the same protocol and column from the RNeasy kit. Quality and concentration of samples was determined using Nanodrop ND-1000 and Qubit readings with the RNA BR Assay kit. RNA samples were sequenced by Novogene (Cambridge, UK) on an Illumina NovaSeq 6000 using a 150 bp paired-end protocol. Gene expression was quantified using Salmon v1.10.2 (Patro et al., 2017) by pseudoaligning reads to CDS from NC_009641.1 (Baba et al., 2008). The resulting counts were modelled as a function of strain using DESeq2 1.40.2 (Love et al., 2014). Differential gene expression was determined by a threshold of absolute fold change greater than 2 and an FDR-adjusted *p*-value less than 0.05. Raw sequencing data has been deposited under BioProject accession number PRJNA1196949.

### Production of recombinant Prs and enzyme activity assay

The gene encoding for wildtype Prs was amplified from Newman genomic DNA using PrimerSTAR PCR mastermix (TaKaRa Bio) and primers listed in Table S2 and cloned into pET28a between the *Nde*I and *Bam*HI sites using In-Fusion Snap Assembly mastermix. The E112K mutation was introduced *via* site-directed mutagenesis and confirmed by bidirectional sequencing. Wildtype and mutant Prs were expressed in *E. coli* BL21 in LB broth with IPTG induction at 37°C for 4 hours. Purification of recombinant Prs was adapted from Walter et al., 2020 and Zhou et al., 2019. Briefly, cultures were pelleted, resuspended in lysis buffer (50 mM potassium phosphate pH 7.6, 300 mM NaCl, 10 mM imidazole, 10% glycerol) containing lysozyme, DNase I and 10 mM MgCl_2_, and lysed on ice by sonication. Prs proteins were purified from the cleared lysates using a HisTrap HP column with increasing concentrations of imidazole. Protein purity was confirmed to be >90% by SDS-PAGE and proteins were exhaustively dialysed against storage buffer (50 mM potassium phosphate pH 7.5, 200 mM NaCl, 10% glycerol) before use.

Prs activity was determined using an established enzyme-coupled spectrophotometric assay (Braven et al., 1984). Reactions (100 µL) contained 5 mM ATP, 5 mM ribose-5-phosphate, 5 mM phosphoenolpyruvate, 1 mM NADH, 10 units myokinase, 7 units pyruvate kinase and 10 units lactate dehydrogenase in assay buffer (50 mM sodium phosphate pH 7.5, 100 mM NaCl, 5 mM MgCl_2_). The mixture was preheated to 37°C and the reaction initiated with the addition of Prs to 1 µM. The oxidation of NADH to NAD^+^ was monitored continuously at 340 nm. The rate of activity was found from the slope over the first 400 seconds and adjusted for background (change in absorbance at 340 nm in the absence of Prs). Each enzyme was tested in quadruplicate.

## Acknowledgments

This work was supported by a project grant from the Canadian Institutes of Health Research (CIHR) awarded to A. B. B. and J. K. H (PJT173349), and a Springboard Award from the Academy of Medical Sciences awarded to J. K. H. A. T. D. was supported by a CIHR Doctoral Foreign Study Award. The authors acknowledge Research Computing at the James Hutton Institute for providing computational resources and technical support for the “UK’s Crop Diversity Bioinformatics HPC” (BBSRC grants BB/S019669/1 and BB/X019683/1), use of which has contributed to the results reported within this paper.

