## Supplemental Material for "*Staphylococcus aureus* COL: An Atypical Model Strain of MRSA that Exhibits Slow Growth and Antibiotic Tolerance Due to a Mutation in PRPP Synthetase"

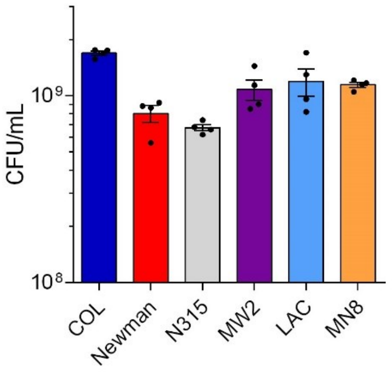


**Figure S1. Stationary phase culture density of COL and other model strains of *S. aureus*.** Four cultures of each strain were inoculated with a single colony of the strain shown, incubated for 18 hours and plated out for viable counting. Data shown are the mean of four biological replicates; error bars (where visible) represent the SEM.

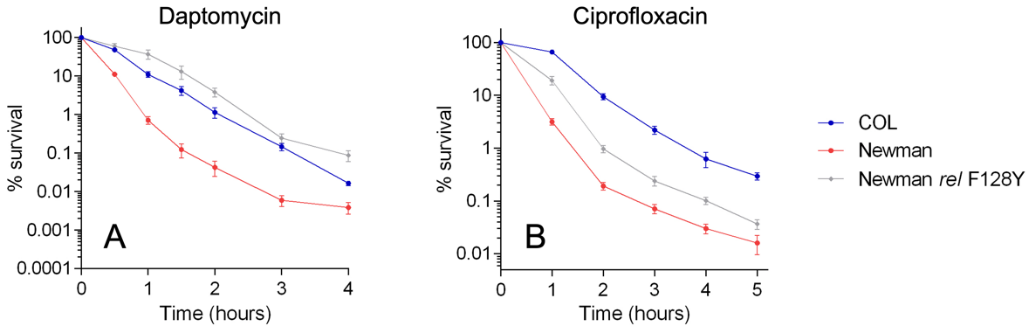


**Figure S2. Comparison of the COL tolerance phenotype to that of a previously characterized tolerant mutant.** Strains were exposed to (A) daptomycin or (B) ciprofloxacin and viable counts were determined at intervals. Counts at each time point are expressed as a percentage of the starting inoculum. Data shown are the mean of three biological replicates; error bars, where visible, represent the SEM.

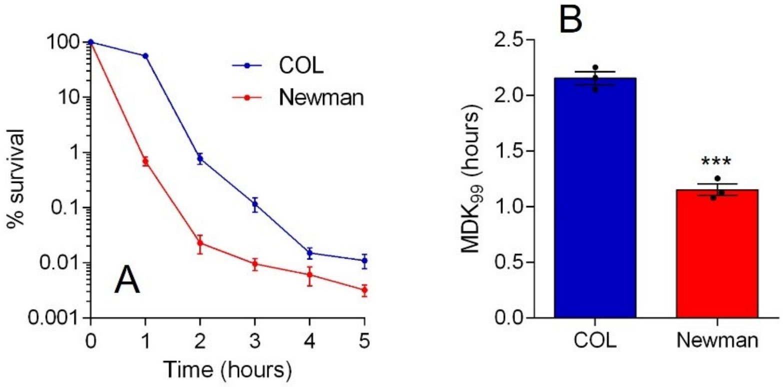


**Figure S3. Exponentially-growing cultures of COL exhibit tolerance to ciprofloxacin.** (A) Cultures of Newman and COL were grown to mid-exponential phase before being diluted to ~$10^6$ CFU/mL, exposed to ciprofloxacin, and viable counts determined at intervals. Counts at each time point are expressed as a percentage of the starting inoculum. Data shown are the mean of three biological replicates; error bars, where visible, represent the SEM. (B) Minimum duration for killing for 99% of the population ($MDK_{99}$) values were interpolated from the data shown in panel A. Asterisks above the Newman bar indicate a statically significant difference between the means as determined by an unpaired *t*-test ($P \leq 0.001$).

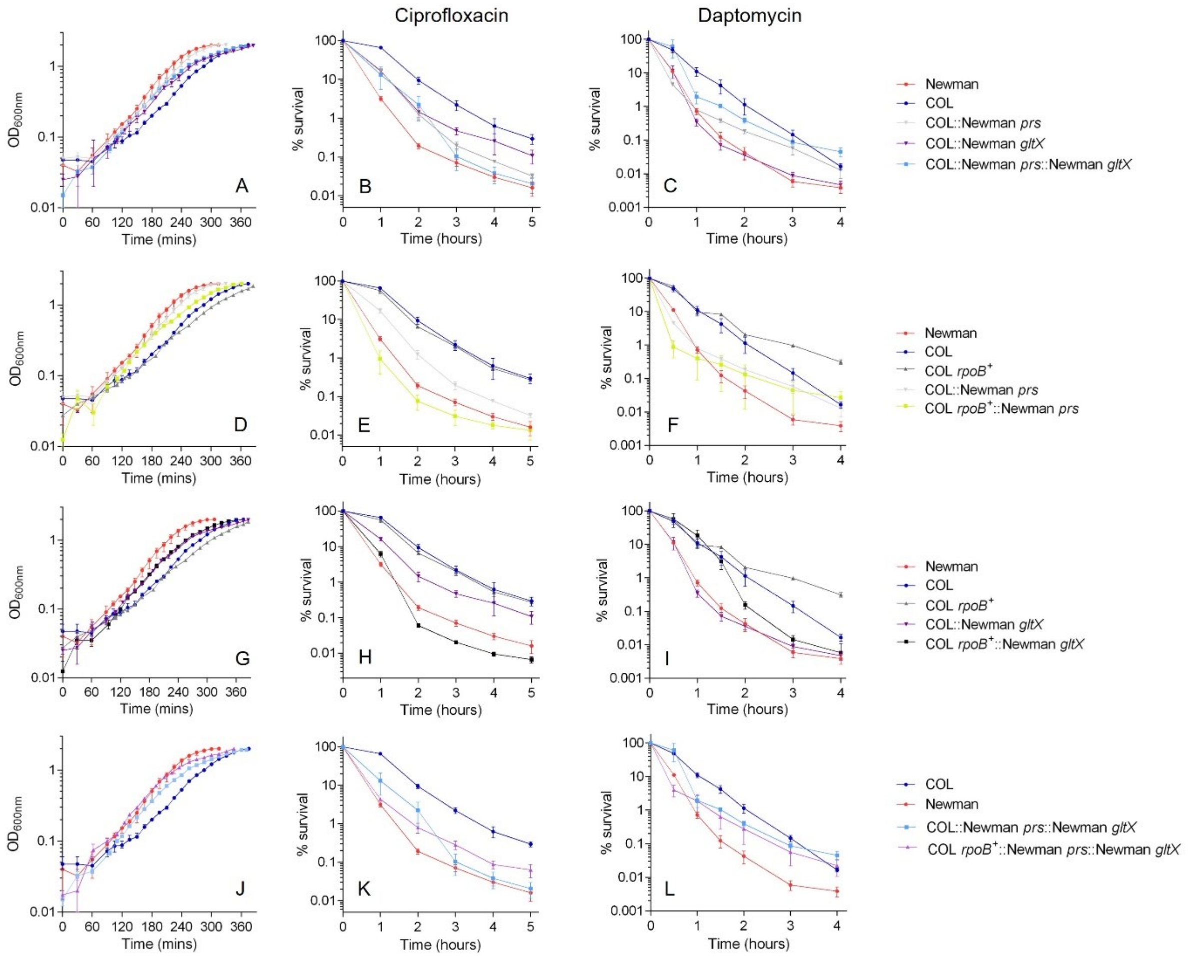


**Figure S4. Impact of double and triple allele swapping on the growth and tolerance of COL.** (A, D, G and J) Growth curves of wildtype COL, wildtype Newman and allele swapped mutants grown in TSB. Data shown are the mean of four biological replicates each derived from a different colony (error bars represent the range). (B, E, H and K) Time-kills with ciprofloxacin. (C, I, F and L) Time-kills with daptomycin. Data shown are the mean of three biological replicates; error bars (where visible) represent the SEM. See Table 2 for lag times, doubling times, and $MDK_{99}$ values derived from these data.

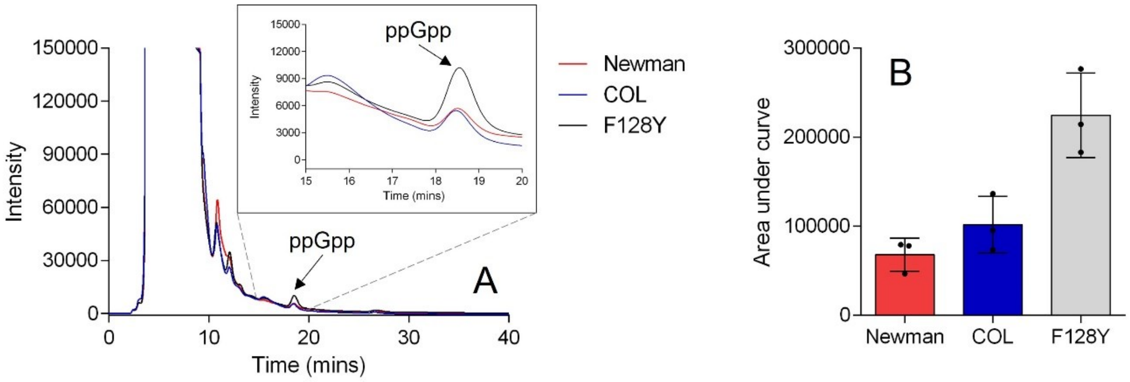


**Figure S5. HPLC quantitation of ppGpp.** (A) Representative HPLC chromatograms for Newman, COL and F128Y overlaid with a ppGpp standard. (B) Area under the curve data for the ppGpp peak. Data shown are the mean of three biological replicates; error bars represent the SEM. Asterisks above bars indicate statically significant differences between means when compared with COL, as determined by a one-way ANOVA with Dunnett's multiple comparisons test (**, indicates *P* ≤0.001; ns indicates not significant).

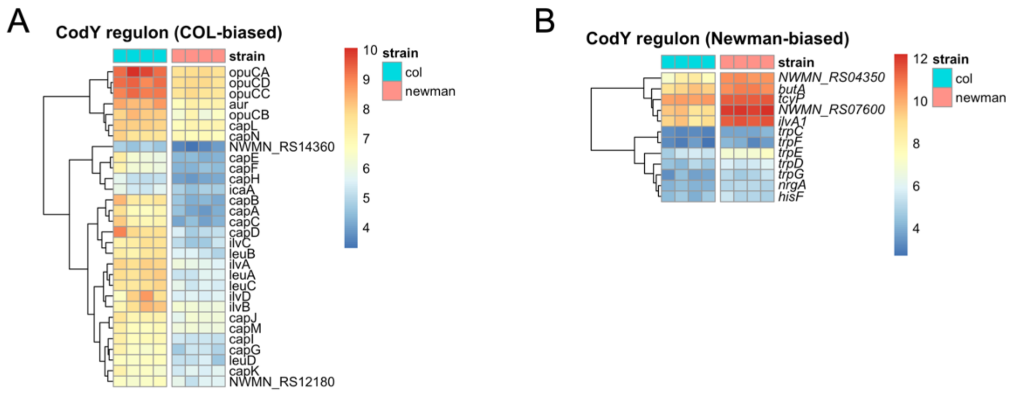


**Figure S6. Differential expression of stringent response-associated genes between COL and Newman.** (A) Heatmap of CodY regulon genes that are significantly upregulated in COL compared with Newman (fold change >2, FDR-adjusted *P*≤0.05). (B) Heatmap of CodY regulon genes that are significantly upregulated in Newman compared with COL (fold change >2, FDR-adjusted *P*≤0.05).

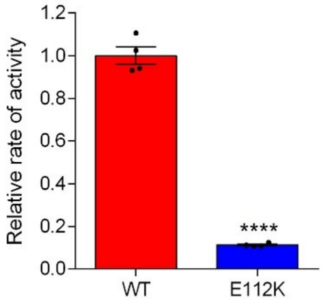


**Figure S7. Impact of E112K mutation on Prs activity.** Relative rate of activity of E112K mutant of *S. aureus* Prs compared with the wildtype enzyme. Activity was measured in a continuous enzyme-coupled assay and the initial rate found from the slope. Bars represent the mean of four replicates; error bars (where visible) represent the SEM. Asterisks above the mutant bar indicate a statistically significant difference from the wildtype mean determined by an unpaired *t*-test (**** indicates *P* ≤0.0001).

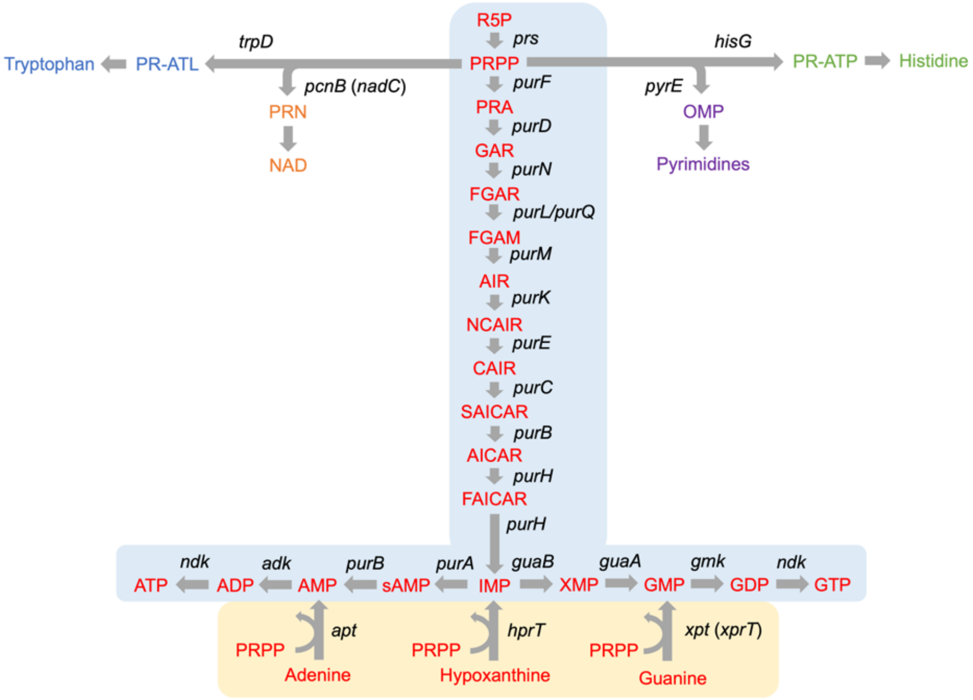


**Figure S8. Metabolic pathways that utilize PRPP.** Six metabolic pathways use PRPP as a substrate for synthesis of important molecules. The *de novo* purine synthesis pathway (shaded in blue) is shown in detail, from ribose-5-phosphate to ATP and GTP *via* inosine monophosphate. The purine salvage pathway (shaded in yellow) also uses PRPP, as does the synthesis of tryptophan, histidine, pyrimidine and NAD. For these pathways, only the enzymes that use PRPP as a substrate are shown. Abbreviations: ribose-5-phosphate (R5P), phosphoribosyl pyrophosphate (PRPP), inosine monophosphate (IMP), oritidine 5'-monophosphate (OMP), 5'-phosphoribosylnicotinate (PRN), phosphoribosyl-ATP (PR-ATP), phosphoribosyl-anthranilate (PR-ATL).

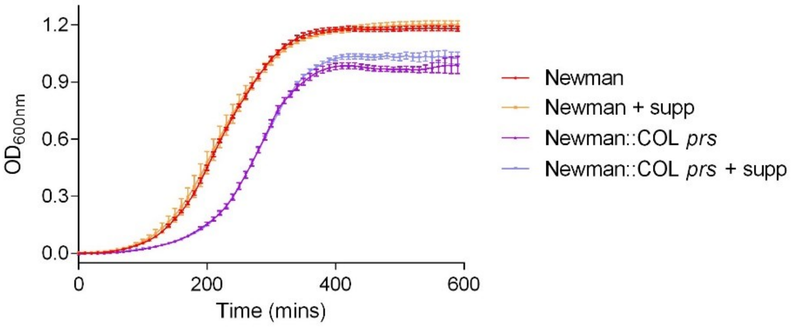


**Figure S9. Effect of media supplementation on the growth of Newman::COL *prs*.** The growth profiles of wildtype Newman and Newman::COL *prs* were compared in TSB and TSB supplemented with histidine, tryptophan, adenosine, guanosine and inosine. Data shown are the mean of six biological replicates; error bars (where visible) represent the SEM.

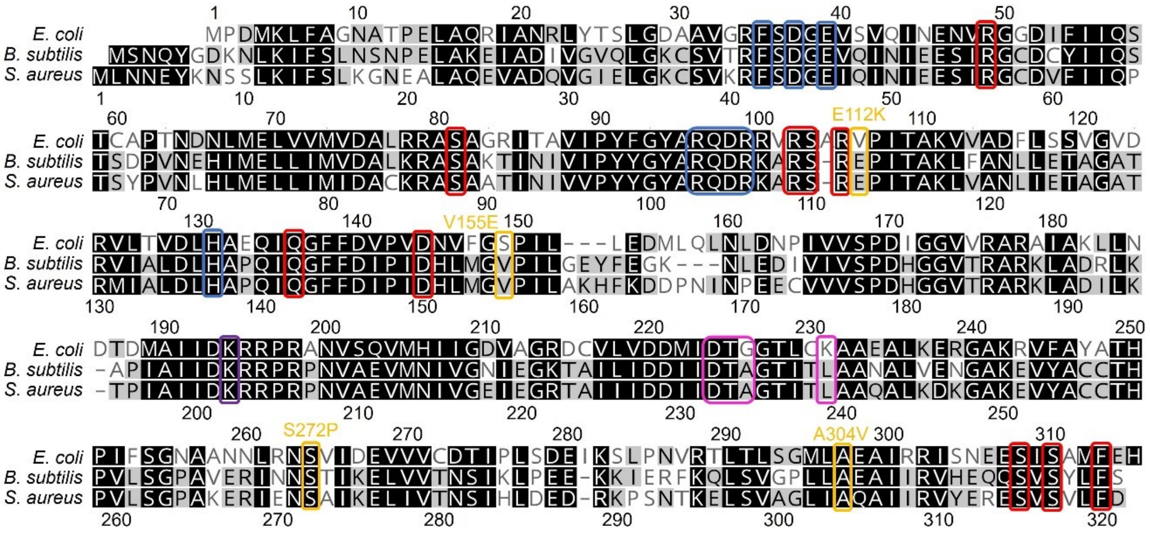


**Figure S10. Alignment of Prs from *E. coli, B. subtilis* and *S. aureus* showing important residues in relation to mutation sites.** Amino acid numbering above and below refer to the *E. coli* and *S. aureus* sequences, respectively. Coloured boxes indicate ATP-binding residues (blue), ribose-5-phosphate-binding residues (pink), allosteric binding site residues (red), and a residue required for essential intramolecular interaction (purple) based on structural and mutational studies. Yellow boxes indicate the Prs mutations identified in COL and clinical isolates from the BSAC collection.
